# Loss of AMPK potentiates inflammation by activating the inflammasome after traumatic brain injury in mice

**DOI:** 10.1101/2024.06.25.600422

**Authors:** Mohammad Ejaz Ahmed, Hamid Suhail, Mohammad Nematullah, Md Nasrul Hoda, Shailendra Giri, Abdullah Shafique Ahmad

## Abstract

Traumatic brain injury (TBI) is a significant public health concern characterized by a complex cascade of cellular events. TBI induces adenosine monophosphate-activated protein kinase (AMPK) dysfunction impairs energy balance activates inflammatory cytokines and leads to neuronal damage. AMPK is a key regulator of cellular energy homeostasis during inflammatory responses. Recent research has revealed its key role in modulating the inflammatory process in TBI. Following TBI the activation of AMPK can influence various important pathways and mechanisms including metabolic pathways and inflammatory signaling. Our study investigated the effects of post-TBI loss of AMPK function on functional outcomes inflammasome activation, and inflammatory cytokine production. Male C57BL/6 adult wild-type (WT) and AMPK knockout (AMPK-KO) mice were subjected to a controlled cortical impact (CCI) model of TBI or sham surgery. The mice were tested for behavioral impairment at 24 h post-TBI thereafter, mice were anesthetized, and their brains were quickly removed for histological and biochemical evaluation. *In vitro* we investigated inflammasome activation in mixed glial cells stimulated with lipopolysaccharides+ Interferon-gamma (LI) (0.1 µg/20 ng/ml LPS/IFNg) for 6 h to induce an inflammatory response. Estimating the nucleotide-binding domain, leucine-rich–containing family pyrin domain containing western blotting ELISA and qRT-PCR performed 3 (NLRP3) inflammasome activation and cytokine production. Our findings suggest that TBI leads to reduced AMPK phosphorylation in WT mice and that the loss of AMPK correlates with worsened behavioral deficits at 24 h post-TBI in AMPK-KO mice as compared to WT mice. Moreover compared with the WT mice AMPK-KO mice exhibit exacerbated NLRP3 inflammasome activation and increased expression of proinflammatory mediators such as IL-1b IL-6 TNF-a iNOS and Cox 2. These results align with the *in vitro* studies using brain glial cells under inflammatory conditions, demonstrating greater activation of inflammasome components in AMPK-KO mice than in WT mice. Our results highlighted the critical role of AMPK in TBI outcomes. We found that the absence of AMPK worsens behavioral deficits and heightens inflammasome-mediated inflammation thereby exacerbating brain injury after TBI. Restoring AMPK activity after TBI could be a promising therapeutic approach for alleviating TBI-related damage.

## Introduction

Traumatic brain injury (TBI) is a leading cause of death and disability worldwide. Patients with TBI and their families often face emotional and financial burdens, requiring long-term care ^1,2^. However, diagnosing and treating TBI patients remains challenging. Current diagnostic methods are not very effective and can be complicated when trying to understand patients’ conditions and outcomes. Finding new targets for treatment could help reduce both the number of people affected and the severity of injuries ^3^.

Both preclinical and clinical studies have shown that TBI leads to a long-lasting reduction in glucose metabolism, which adversely affects brain function. ^4–6^. This means that the brain’s ability to use glucose for energy is impaired after TBI, impacting brain function. ^7^. AMPK is a serine/threonine kinase that acts as a master regulator of cellular energy balance and governs diverse cellular processes crucial for maintaining cellular homeostasis. AMPK plays a major role as a cellular energy sensor and is activated in response to low-energy conditions, such as ATP or an elevated AMP/ATP ratio ^8,9^. Post-TBI, AMPK activation typically restores the cellular energy balance rather than suppressing energy consumption and ATP upregulation. ^10–12^. AMPK is expressed in various cell types within the central nervous system, including neurons, astrocytes, oligodendrocytes, and microglia. The activation of AMPK in these different cell types contributes to a range of cellular responses critical for maintaining brain homeostasis and responding to injury and inflammation ^13,14^. In the brain, AMPK is activated in response to various pathological conditions, including hypoxia, excitotoxicity, and increased AMP/ATP or ADP/ATP ratios ^15–17^. AMPK activation serves as a critical cellular response to maintain energy homeostasis and protect against metabolic stress in the brain.

Recently, AMPK and the inflammasome have attracted the attention of clinical and preclinical researchers. The involvement of AMPK in the progression and pathogenesis of several diseases has led to the development of new therapeutic strategies. AMPK has emerged as a key player in modulating inflammation, particularly through its interaction with the inflammasome. Inflammasome is a multiprotein complex composed of three protein subunits: a sensor molecule, NLRP3; an adaptor protein, ASC; and an effector protein, caspase 1. Inflammasome primes the natural defense system of the body and subsequently triggers the activation of the inflammatory response by promoting the maturation and secretion of proinflammatory cytokines, such as interleukin-1β (IL-1β) and interleukin (IL-18) ^18–21^

Many studies have shown that TBI elicits a complex immune response characterized by the activation of NLRP3 inflammasome components ^22,23^. One of the initial studies conducted by Liu et al. on inflammasome showed that most inflammasome markers were upregulated 6 h after TBI and remained upregulated until 7 days after the injury ^24^. Other studies have shown that NLRP3, caspase-1, and IL-1β expression is significantly increased at 12 h and 72 h post-TBI in mice subjected to cold brain injury ^25,26^. Among all inflammasomes in the brain, NLRP3 facilitates the cleavage of caspase-1 and interleukin-1β (IL-1β), which potentiate the inflammatory response against TBI ^27–29^. In an animal model of TBI, the levels of NLRP3 and the inflammasome complex increased in the peri-lesion area of the cortex and were correlated with long-term functional outcomes after TBI ^30^.

The present study revealed an important role of AMPK/NLRP3 inflammasome-mediated neuroinflammation in TBI. To our knowledge, no direct studies have investigated the role of the loss of AMPK and NLRP3 inflammasome activation in the CCI-induced model of TBI.

## Materials and Methods

### Animals and experimental groups

Male C57BL/6 adult wild-type (WT) and AMPK-KO mice were maintained and housed at the animal house of Henry Ford Health. Mice were randomly divided into four groups: (i) WT sham group (n=10 mice per group), (ii) WT+TBI group (n=10 mice per group), (iii) AMPK-KO sham group (n=10 mice per group), and (iv) AMPK-KO+TBI group (n=10 mice per group). All the experiments were conducted according to the protocol approved by the Henry Ford Health Institutional Animal Care and Use Committee (IACUC). Mice were kept at room temperature (21 ± 1°C) and 50 ± 5% humidity on a 12 h dark/light cycle. All the mice were given access to food, water, and ad libitum.

### Induction of traumatic brain injury

Male C57BL/6 adult WT and AMPKα-KO mice underwent either sham or TBI surgery using a controlled cortical impact (CCI) method, which has been described previously ^31^. Briefly, mice were anesthetized with 3% isoflurane and were kept on a stereotaxic frame and maintained with 1.5%-2% isoflurane throughout the surgical procedure using a SomnoSuite low-flow anesthesia system (Kent Scientific Corporation). A craniotomy was performed in the right parietal bone midway between the bregma and lambda using a hand twist drill and coordinated 1 mm lateral to the midline, leaving the dura intact. The mice were impacted at 3 m/s with an 85 ms dwell time and 3.0 mm depression using a 3 mm diameter tip (Pin Point PCI3000 Precision Cortical Impactor, Hatters Instruments). Sham-operated mice underwent the same surgical procedure, but there was no impact on the skull. Body temperature was maintained at 37.0 ± 0.5°C throughout the procedure using small-animal temperature control (Kopf Instrument). The mice were returned to a clean home cage that was placed on an automatic warming pad (Kent Scientific Corporation).

### Behavioral analysis

#### Open field test

The open-field test is used to evaluate rodent locomotor and exploratory activities in rodents ^32,33^. The open field apparatus consists of a plastic box with a floor base dimension of 96 cm × 96 cm and a wall dimension of 50 cm. White lines (6 mm) are drawn on the base to create 16 identical squares. Each mouse was positioned in the middle of the open field apparatus and was subjected to a 5-minute observation period. The amount of time spent by individual mouse in the center zone was recorded and analyzed by using automated ANY-maze video tracking software. The open field apparatus was cleaned with 70% ethanol between each test.

#### Elevated plus maze test

The elevated plus maze (EPM) test is the most widely used method for assessing anxiety-like behavior in rodents in a novel environment ^34^. The maze consisted of two opposing open arms (50 cm × 10 cm × 0.5 cm) and two opposing closed arms (50 cm × 10 cm × 40 cm), which were crossed perpendicularly with an open center area (10 cm × 10 cm) elevated 50 cm from the floor ^35^. The EPM test was performed at 24 h post-TBI. Individual mice were placed in the center of the EPM and subjected to 5 minutes of observation. The activity of the mice was recorded by automated ANY-maze video tracking software. Measures of anxiety were assessed by measuring the time spent in the open arm and the number of times the participants entered the open arm. Mice with greater anxiety will spend less time in the open arm. The EPM apparatus was cleaned with 70% ethanol between each test.

#### Rotarod test

Motor deficits test in post-TBI mice were assessed by using a rotarod device, consist of 30 mm in diameter rotating, rough solid surface rods that are suspended from the floor at a height of 30 cm and divided into 4 separate compartments to conduct four mice tests simultaneously. Twenty-four hours before TBI induction, the mice were acclimatized for 3 min on the rotarod with accelerating speed reaching a maximum speed of 20 RPM at 3 min ^36,37^. Each mouse was given 3 min (180 sec) and the speed of the rotarod was set at 20 RPM for 3 trials. About 10 min intervals were given between each trial for the mouse to rest. The average of three trials was used to calculate the average retention time for each mouse on the rotarod.

#### Hanging wire test

Grip strength was assessed at 24 h post-TBI in mice as previously described ^33^. To assess grip strength, a wire was stretched between two poles 100 cm apart and 100 cm above the floor. Briefly, the mouse was held with its tail and allowed to grasp the wire with its front paws then the support was withdrawn, and the mouse was allowed to hang from the wire with both forepaws. Each trial was evaluated on a 5-point scale: 0, fall off; 1, hang onto the wire by both forepaws; 2, hang onto the wire by both forepaws with attempts to climb onto the wire; 3, hang onto the wire by both forepaws as well as one or both hind limbs; 4, hang onto the wire by forepaws with the tail wrapped around the wire; and 5, escape by reaching to the role. The highest reading of three successive trials was taken for each mouse ^38^. To prevent any injury to the mouse after the fall, a thick, soft mat was placed on the floor.

### Western blot analysis

Western blot analysis was performed as previously described ^39^. The mice were anesthetized with isoflurane and decapitated at 24 h post-TBI. Thereafter, the brains were quickly removed, and the ipsilateral part of the brain was microdissected and cryopreserved in liquid nitrogen. Briefly, 35 µg protein samples from each mouse were equally loaded into each well and separated on 4–20% SDS PAGE gels, then, protein bands were transferred to nitrocellulose transfer membranes (Thermo Scientific) and blocked with 5% nonfat milk for 1 h. Then, the membranes were incubated with the appropriate primary antibodies against AMPKα (1:1000 BS 10344R, Bioss, Woburn, MA), pAMPKα (Thr183/Thr172; 2535S, Cell Signaling), and the NLRP3 (15101, Cell Signaling) overnight at 4°C. After three washes with TBST, the membranes were incubated with the appropriate HRP-conjugated secondary antibody for 2 h at room temperature. Followed by membranes were washed, and the protein bands were visualized with an enhanced chemiluminescence (ECL) solution (Clarity Western ECL Substrate, 1705062 Bio-Rad Labs, Des Plaines, IL). Quantitative analysis of the bands was performed with ImageJ software (version 1.49; NIH).

### Cresyl violet staining and quantification of lesion size

Coronal sections (25 µm thick) of mouse brains were prepared using a cryostat (Thermo Fisher Scientific, MA, USA). Sections were collected for lesion size quantification. Every fifth section was selected and stained with 0.1% (*w/v*) cresyl violet for 5 min. The sections were then dehydrated with a graded concentration of ethanol followed by xylene. The mean lesion size was quantified by using ImageJ software (NIH). The lesion size was calculated by subtracting the area of the ipsilateral hemisphere from the total area of the contralateral hemisphere. ^38,40^.

### Immunofluorescence staining and quantification

Mice were anesthetized with isoflurane and transcardially perfused with 0.9% saline followed by 4% paraformaldehyde (PFA) at 24 h after TBI. Brains were harvested and processed for immunofluorescence staining as previously described ^39,41^. Using a cryostat (Thermo Fisher Scientific, MA, USA), 25–µm sections were cut on a cryostat. Following washing, permeabilization, and blocking with 0.4% PBST for one hour at room temperature, the sections were incubated with primary antibodies against NeuN (PA5-78499, Invitrogen), MAP2 (MAB3418, Millipore), GFAP (PA5-16291, Thermo Fisher), and Iba1 (016-20001, Wako) at 4°C overnight. The next day, the sections were washed 3 times with PBST and incubated with Alexa Fluor 568/488-conjugated goat anti-mouse/rabbit secondary antibody (1:500; Thermo Scientific Rockford, IL) for 1 h at room temperature. Finally, sections were washed and mounted on slides using a Vectashield mounting medium with 4′,6-diamidino-2-phenyl indole (DAPI) (H-1200; Vector Laboratories, Inc., CA, USA) and a coverslip. Confocal images were obtained using Zeiss AxioTvert 200/Axiovert 200 M microscopes. Using ImageJ software (NIH), five randomly chosen fields from each animal’s cortex were quantified in four nonadjacent sections spaced approximately 100 µm apart.

### Primary brain glial cell culture

The brains of 2-3-day-old WT and AMPK-KO rat pups were used to prepare mixed glial cell cultures. We followed our previously published protocol to prepare primary mixed glial cell cultures ^42–45^. In brief, the brain was removed, and cortical cells were separated from blood vessels and the meninges. The cerebral cortical cells were mechanically triturated into a single-cell suspension with a Pasteur pipet and suspended in complete media (DMEM 4.5 g/L) supplemented with 10% FBS and 10 mg/mL antibiotics. The suspended cells were seeded in 75 cm^2^ poly-D-lysine-coated flasks and incubated at 37°C in a humidified 5% CO2 incubator. After 24 h, the media was replaced with fresh complete DMEM (4.5 g/L), which was then exchanged twice a week. The cells were grown for at least 12-14 days until they reached full confluence and were ready to dissociate.

### Brain Glial Cell Stimulation

Glia isolated from WT and AMPK-KO mice were isolated and seeded at a density of 50 × 10^4^ cells per well in 24 well plates containing DMEM supplemented with 10% fetal bovine serum (FBS). The cells were cultured overnight at 37°C in a humidified atmosphere with 5% CO2. The following day, the cells were serum-starved by replacing the culture medium with serum-free DMEM for 2 h after serum starvation, the cells were stimulated with LI (0.1 µg/20 ng/ml LPS/IFNg) to induce an inflammatory response. Cell supernatant was used for cytokines ELISA and cell lysate was processed for qPCR and immunoblot analysis.

### RNA extraction, cDNA synthesis, and quantitative PCR

After stimulation with LI, WT, and AMPK-KO glia were isolated at 6 h, and total RNA was extracted using TRIzol and a RNeasy Kit (Qiagen) according to the manufacturer’s protocol. cDNA was synthesized from 1 µg of total RNA using an iScript reverse transcriptase cDNA synthesis kit (Bio-Rad) to amplify all genes. The primer sets and their nucleotide sequences were selected from the available online database and purchased from Integrated DNA Technology (IDT Coralville, Iowa, USA) or real-time primers. The iTaq Univer SYBR Green Supermix was purchased from Bio-Rad. The thermal cycling amplification conditions were as follows: activation of DNA polymerase was used at 95°C for 3 min, followed by 40 cycles of amplification at 95°C for 30 s and 60°C for 30 s. The expression of the ribosomal protein L-27 housekeeping gene was used to calculate the normalized expression (Ct) of the target gene transcription using CFX Maestro Software.

### Cytokine analysis by enzyme-linked immunosorbent assay (ELISA)

To analyze cytokine levels, isolated glia was stimulated with LI (0.1 µg/20 ng/ml LPS/IFNg). After 6 h of stimulation, the Petri plates were washed three times with warm serum-free media, and the supernatant was collected to measure inflammatory cytokine production via ELISA.

### Statistical analysis

The data are presented as the mean ± SEM and statistical analyses were performed by using GraphPad Prism 9. To determine the significant difference among the groups, Student’s t-test or one-way analysis of variance (ANOVA) followed by Tukey’s post-hoc test was performed. A p < 0.05 was considered to be significant.

## Results

### Effect on the AMPK level in the post-TBI mouse brain

WT mice brains were extracted 24 h after TBI and a western blot was performed to assess the levels of AMPK. Representative western blot images and quantitative data **(Fig. 1A-B)** showed a significant decrease in phosphor-AMPKα1 immunoactivity at 24 h post-TBI compared with sham-operated mice. This suggests that the loss of AMPK activity after TBI affects the total level of the AMPK.

**Figure 1.**
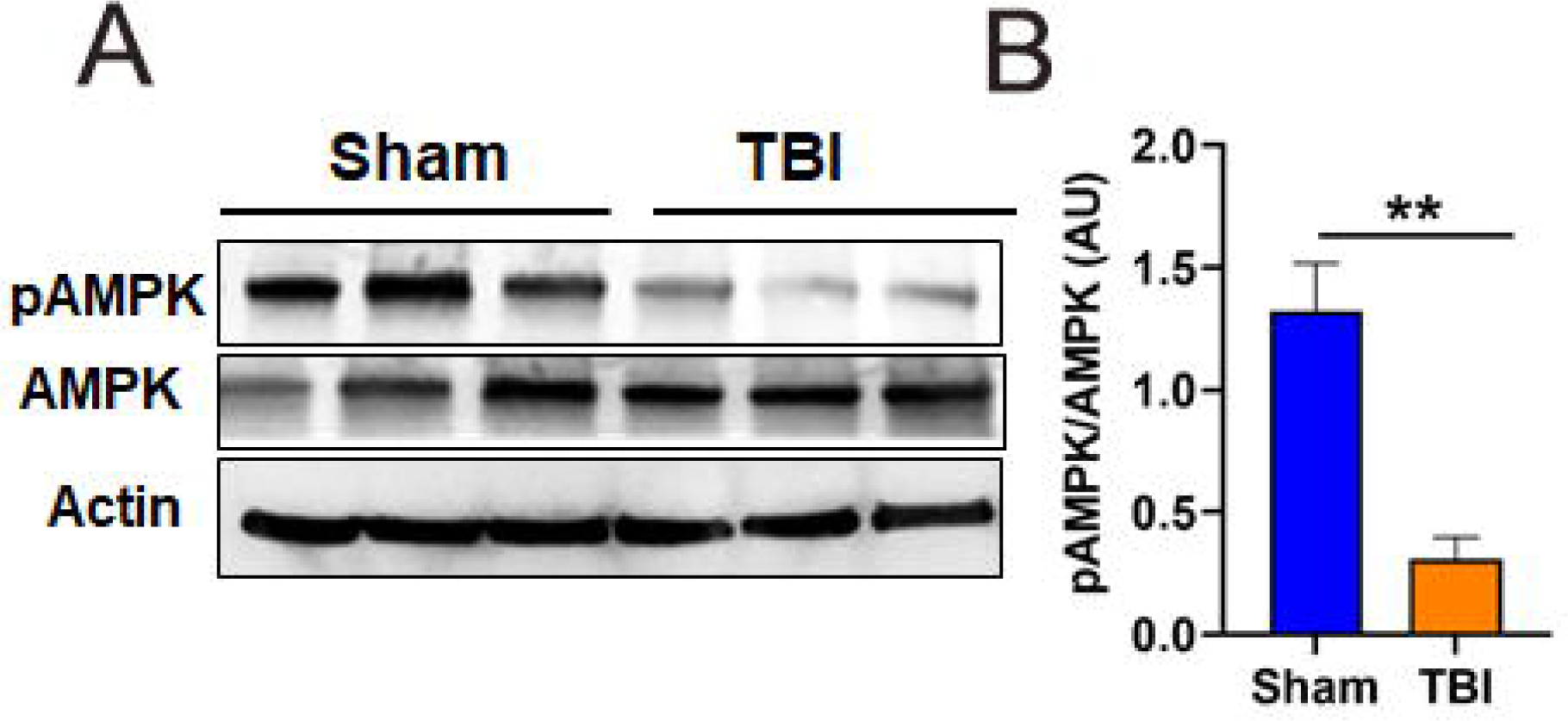
Loss of AMPK phosphorylation following TBI in mice. **(A)** Representative western blot showing the loss of AMPK phosphorylation at 24 h in WT mice. **(B)** Quantitative analysis of AMPK in the WT sham and TBI mice. The data are expressed as the mean ± SEM (n=3/group), **P<0.01.

### AMPK deficiency worsened functional deficits in post-TBI mice

Since AMPKα1 has been reported to be anti-inflammatory ^46–49^, we used global knockout (KO) of AMPKα1 and examined functional deficits in WT and KO mice after TBI. Using the open field apparatus, spontaneous locomotor and anxiety activity were measured by timing how long the mice spent in the center of the field. When the AMPK-KO mice were compared to the WT TBI mice, the AMPK-KO group spent notably less time in the central zone **(Fig. 2A, E)**. To evaluate exploratory and anxiety-like behaviors in mice after TBI, we also used the elevated plus maze test. Compared with WT TBI mice, AMPK-KO mice spent less time in the open arm at 24 h after TBI, indicating that AMPK deficiency promotes anxiety-like behavior **(Fig. 2 B, F)**. The grip strength and motor coordination of the mice were tested by the hanging wire test and rotarod test as shown in **(Fig. 2C, D)**. The results revealed that the AMPK-KO TBI mice exhibited greater deficits than the WT TBI mice. Overall, our results show that the loss of AMPK further aggravates functional deficits after TBI.

**Figure 2.**
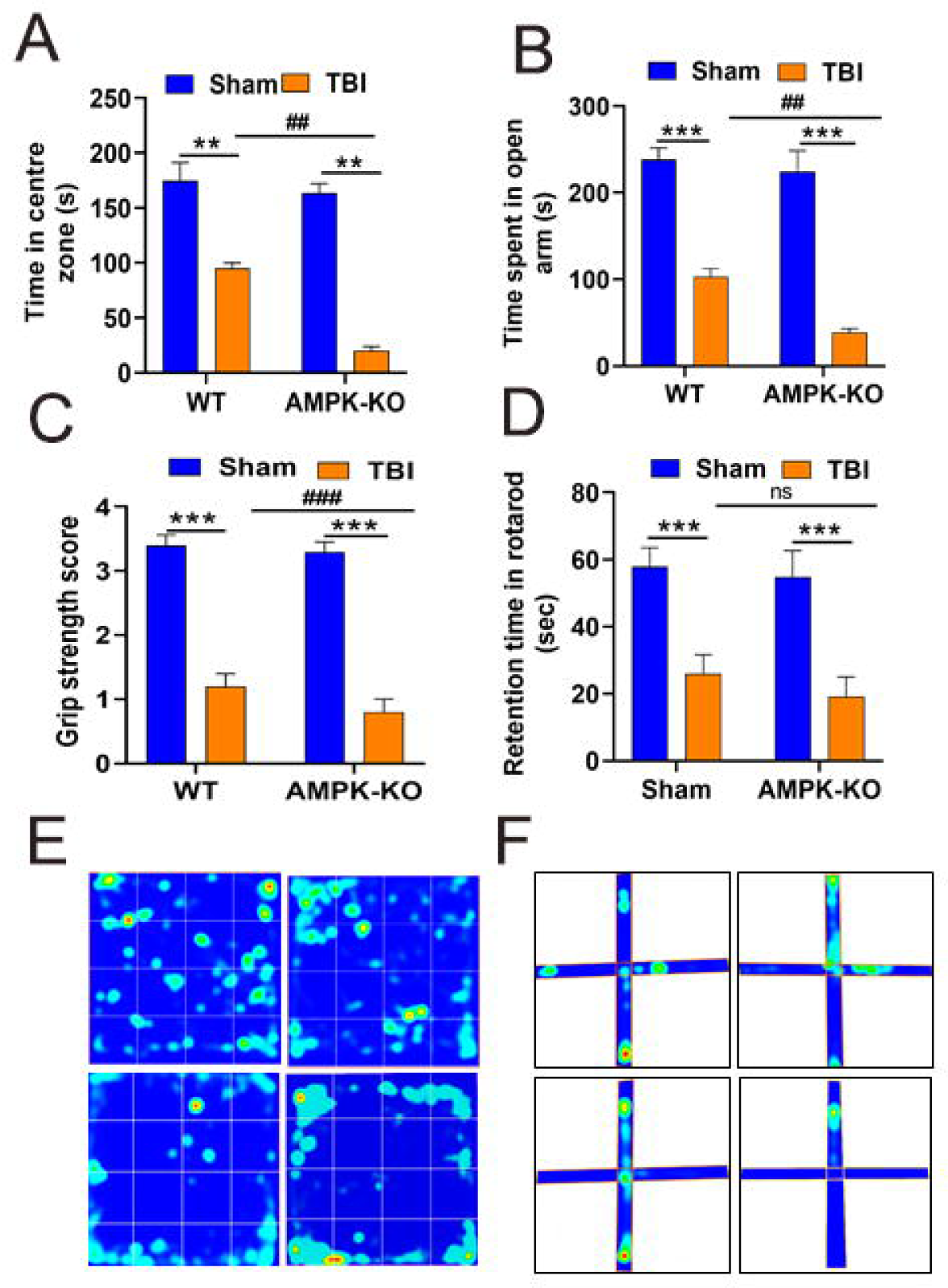
AMPK deficiency worsened functional impairments in TBI mice. The functional impairment was tested at 24 h after TBI**. (A)** The open field test was performed 24 h after TBI. TBI causes mice to spend less time in the central zone (a measure of increased anxiety after TBI. Compared with WT TBI mice, AMPK-KO mice spent significantly less time in the center zone. **(B)** An elevated plus maze test was conducted to measure anxiety-like behavior in the mice. Compared with WT TBI mice, TBI-induced AMPK-KO mice spent significantly less time in the open field test. **(C)** The grip strength test was used to measure dexterity in TBI mice hanging from the suspended wire. Compared with WT mice, AMPK-KO mice exhibited worse scores at 24 h post-TBI. **(D)** Rotarod testing was performed to test motor and coordination activity following TBI in mice. Compared with the WT TBI mice, the AMPK-KO TBI-induced mice stayed on the moving rotarod less often. **(E, F)** Heat map obtained from the open field and elevated plus maze tests showing an increase in the amount of time spent within the center zone (a measure of decreased anxiety). The data are expressed as the means ± SEM (n=10/group), **P<0.01, ***P<0.001, ##P<0.01.

### Loss of AMPK increases lesion size in TBI mice

To investigate the effect of the loss of AMPK on neuronal damage and lesion size in mice 24 h after TBI, WT, and AMPK-KO mice were subjected to unilateral CCI-induced TBI. Representative whole-brain images demonstrating intracranial hemorrhage and lesion size in WT and AMPK-KO mice at 24 h post-TBI (**Fig. 3 A**). We found that intracranial hemorrhage and lesion size were increased in AMPK-KO compared with WT TBI mice. We also evaluated lesion size in the ipsilateral part of the brain We found lesion size was significantly increased in AMPK-KO compared with WT TBI mice **(Fig. 3 B).** These results indicate that AMPK-KO mice exhibit greater tissue damage and bleeding than WT mice.

**Figure 3.**
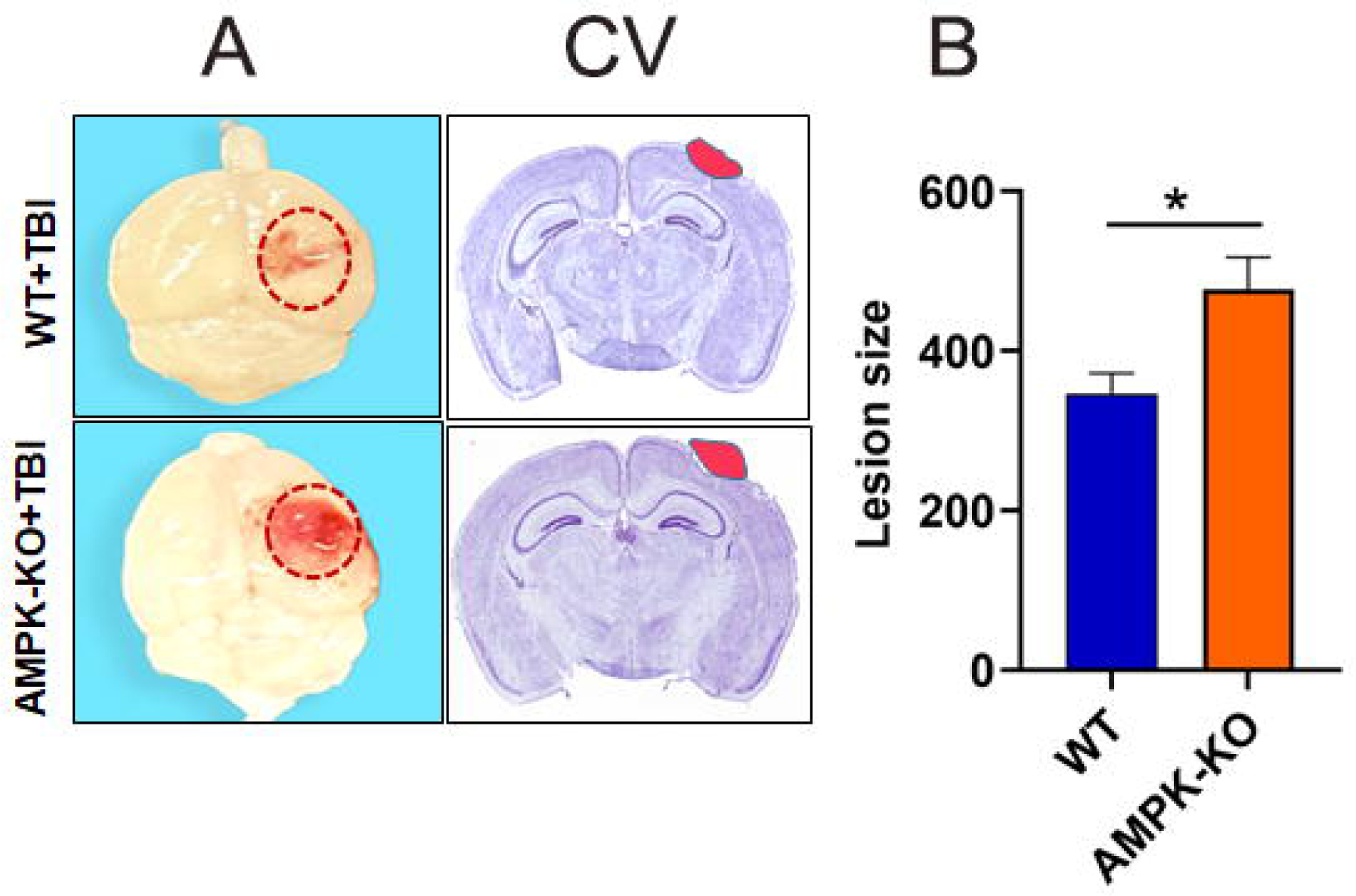
Effect of the absence of AMPK on histological and anatomical changes in TBI mice. (A) Blood clots and tissue damage at the impact site were assessed in WT and AMPK-KO mice at 24 h post-TBI. **(B)** Representative images of cresyl violet staining and quantitative analysis of lesion size are shown. The data are expressed as the mean ± SEM (n=5/group) for each group *P<0.05.

### Loss of AMPK exacerbates TBI-induced neuronal damage in the mouse brain

We further examined the impact of AMPK loss on TBI-induced neuronal loss using immunofluorescence staining. Briefly, brain tissue sections from WT and AMPK-KO mice were immunostained for two neuronal-specific markers, microtubule-associated protein (MAP2) (green) and NeuN (red). MAP2 predominantly stains the cytoskeleton of neurons, such as dendrites, and NeuN is a neuron-specific marker used to indicate neuronal survival following TBI**. (Fig. 4 A, B)**. Compared with WT TBI mice, AMPK-KO TBI mice showed significantly less NeuN staining and poor MAP2 intensity **(Fig 4 C, D),** indicating significant and gradual neuronal loss 24 h post-TBI.

**Figure 4.**
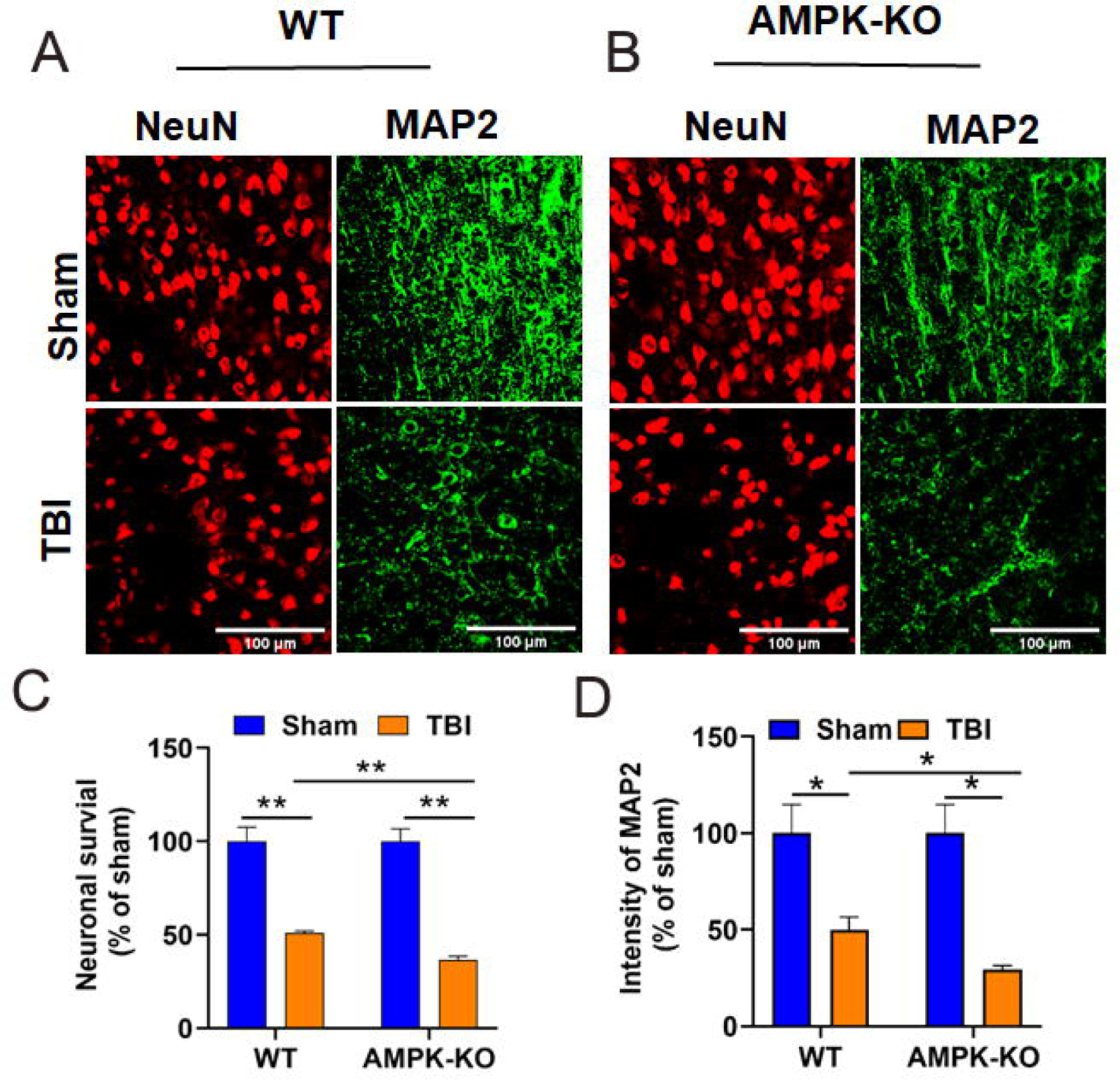
The absence of AMPK causes neuronal damage in TBI mice. **(A, B)** Representative images of immunofluorescence staining of the peri-contusion area of WT and AMPK-KO mice showing MAP2 (green) staining and NeuN (red) staining 24 h following TBI. The sham group of WT and AMPK-KO mice exhibited greater MAP2 intensity with log dendrites. However, the dendrites of the AMPK-KO mice were less intense and degenerated than those of the WT TBI mice. We also stained brain tissue with NeuN to assess neuronal survival at 24 h after TBI. Representative immunofluorescence images showed greater NeuN expression in the WT sham group than in the WT TBI group. We also examined the effect of the loss of AMPK on NeuN expression in AMPK-KO sham and TBI mice. We detected a lower level of NeuN expression in TBI mice than in WT mice. **(C)** Quantitative analysis of MAP2 and surviving neurons. The results are representative of 4-5 sections from each (n=6) mouse per group. Scale bar = 100 μm. *P<0.05, **P<0.01.

### AMPK deficiency exacerbates NLRP3 inflammasome activation in TBI mouse brains

Inflammasomes are essential for combating the inflammatory reaction that occurs after TBI. At 24 h after injury, the brains of the AMPK-KO TBI mice had significantly greater expression levels of NLRP3, caspase-1, IL-1β, and ASC than those of the WT TBI mice, as shown by representative western blot images **(Fig. 5 A, B)**. Quantitative analysis shows significantly increased NLRP3, caspase-1 IL-1β, and ASC in AMPK-KO compared with WT TBI mice **(Fig 5 C, D).** These data indicate that TBI causes the NLRP3 inflammasome to become activated in the injured brain, which is consistent with the findings of other studies.

**Figure 5.**
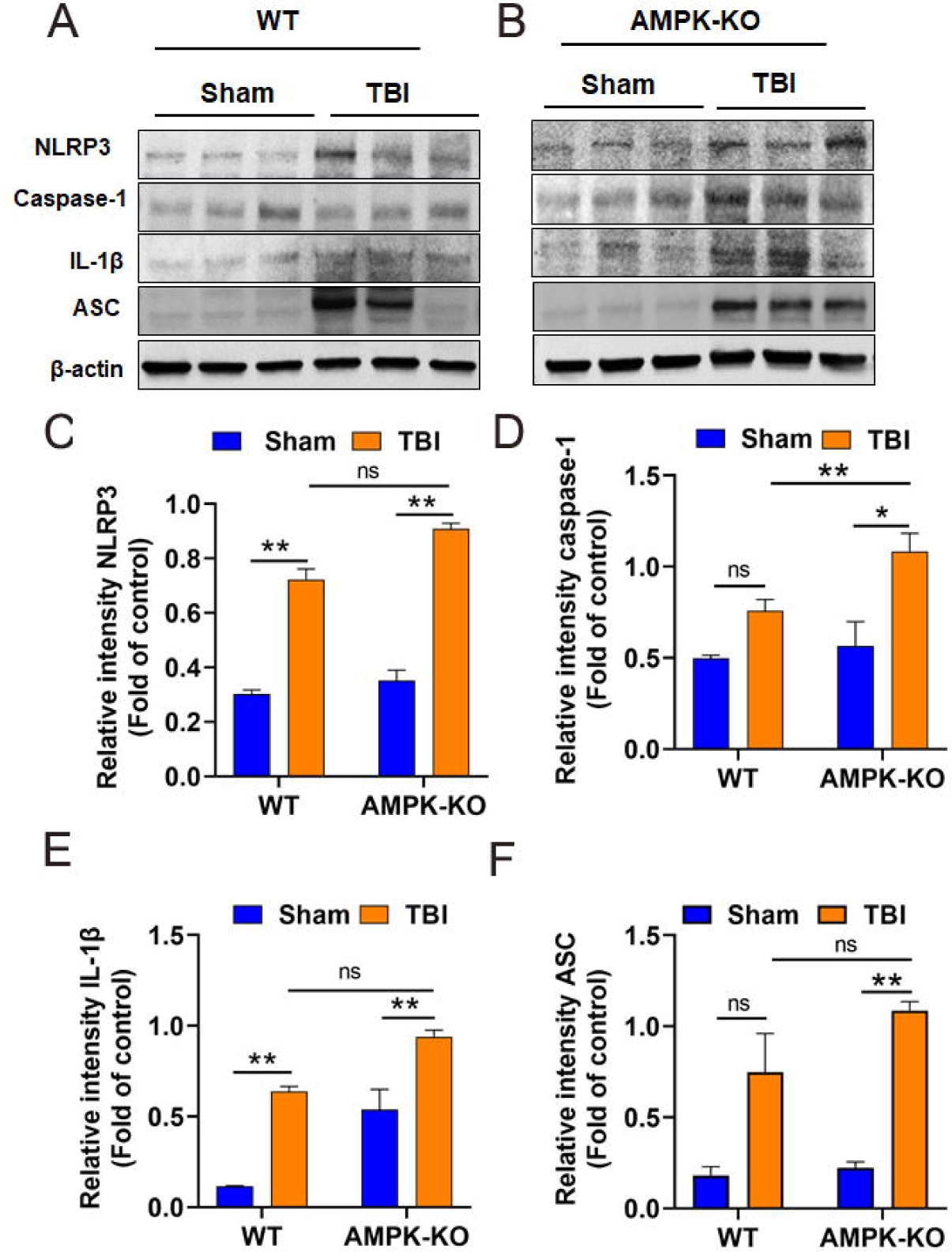
AMPK deficiency potentiates inflammasome activation in TBI mouse brains. Western blot analysis of the inflammasome component markers NLRP3, caspase-1, IL-1β, and ASC revealed that compared with WT TBI mice, AMPK-KO mice exhibited increased expression of inflammasome components following TBI **(A, B)**. **(C-F)** Quantitative analysis of the band intensity normalized to that of actin. The data are presented as the means ± SEMs (n=3/group). *P<0.05, **P< 0.01, ns= not significant.

During TBI, a loss of AMPK and an increase in the level of the NLRP3 inflammasome were observed in WT mice, suggesting a potential inverse relationship between AMPK and the inflammasome. To examine this relationship, we used brain glial cells from WT and AMPKa1 KO pups that were stimulated under inflammatory conditions (0.1 µg/20 ng/ml LPS/IFNg) and examined inflammation and inflammasome status. As depicted in **(Fig. 6 A)**, the production and expression of inflammatory markers, including IL1β, IL6, and TNFα, were significantly greater in the AMPK-KO glial cells than in the WT glial cells under inflammatory conditions. Loss of AMPK also increased the levels of inflammatory mediators, including iNOS and Cox2, compared to those in WT glial cells **(Fig. 6B)**, which was accompanied by increased inflammasome activity, including that of ASC, cleaved caspase 1 and IL1b, further suggesting that AMPK negatively regulates inflammation and the inflammasome under inflammatory conditions.

**Figure 6:**
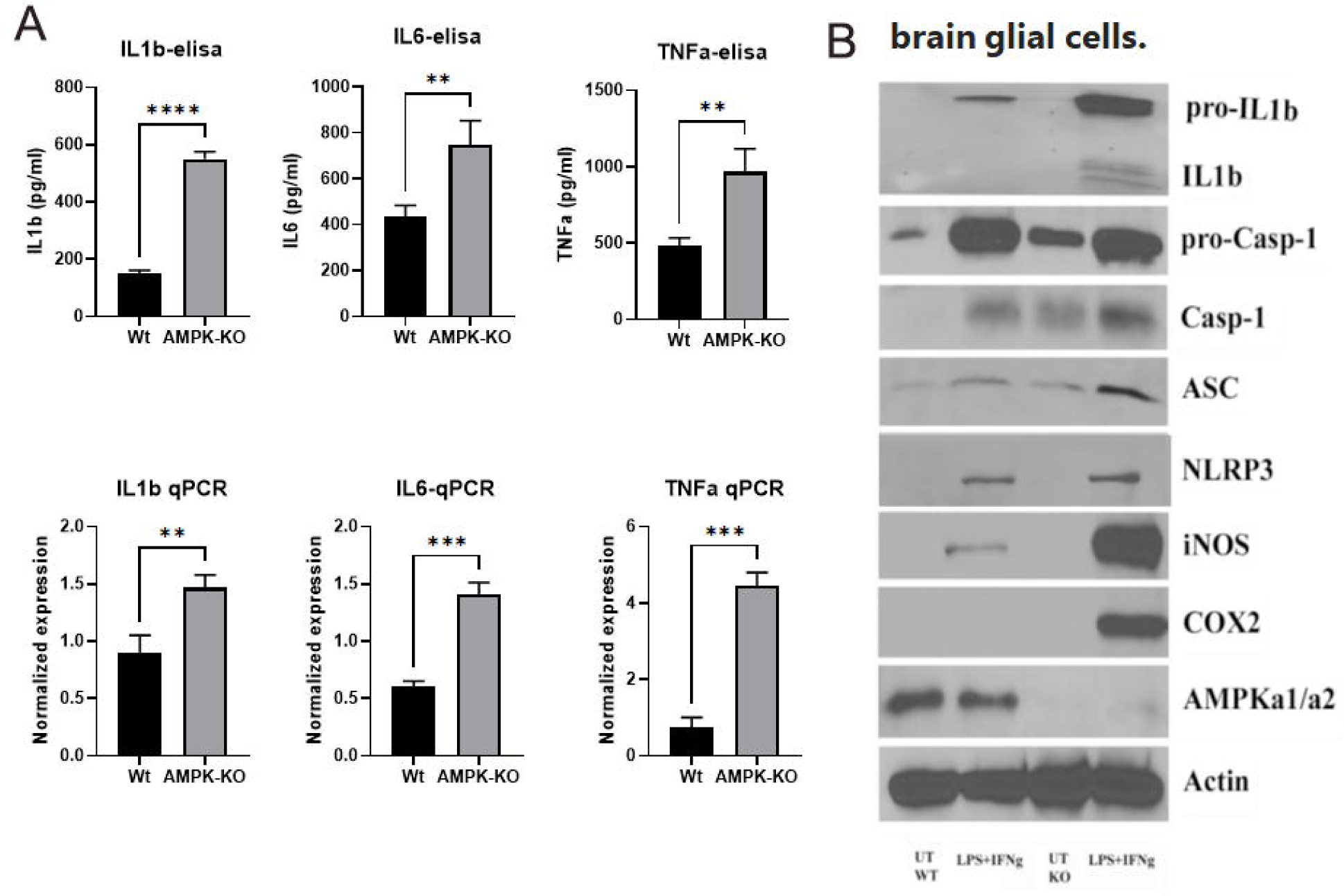
Loss of AMPK in brain glial cells leads to hyperinflammation and inflammasome activation. WT and AMPK-KO brain glial cells were treated with LPS/IFNg (0.1 µg/20 ng/ml) to induce inflammation, the cell supernatant was used for the detection of IL1b, IL6, and TNFa, and the cells were processed for mRNA detection by qPCR (n=4) **(A)**. The cell lysates were processed for immunoblot analysis of various inflammatory mediators and inflammasome proteins using specific antibodies as described in the Methods section **(B)**. Blots are representative of two experiments with similar observations. **P< 0.01, ***P < 0.001, P<0.0001.

### AMPK deficiency increases TBI-induced gliosis in the mouse brain

We investigated how reactive gliosis is impacted by AMPK loss following TBI. Immunofluorescence staining was performed to test microglial and astrocyte activation in the cerebral cortex at 24 h after TBI. Immunostaining shows higher immunoreactivity of GFAP (green), a marker for astrocytes, and Iba1 (red), a marker for microglia in AMPK-KO mice **(Fig 7 A, B)**. Quantitative analysis revealed an increase in GFAP and Iba1 expression in the ipsilateral cortex of the AMPK-KO compared with WT TBI mice, no significant difference was detected between the WT and AMPK-KO TBI mice **(Fig 7 C, D)**

**Fig 7.**
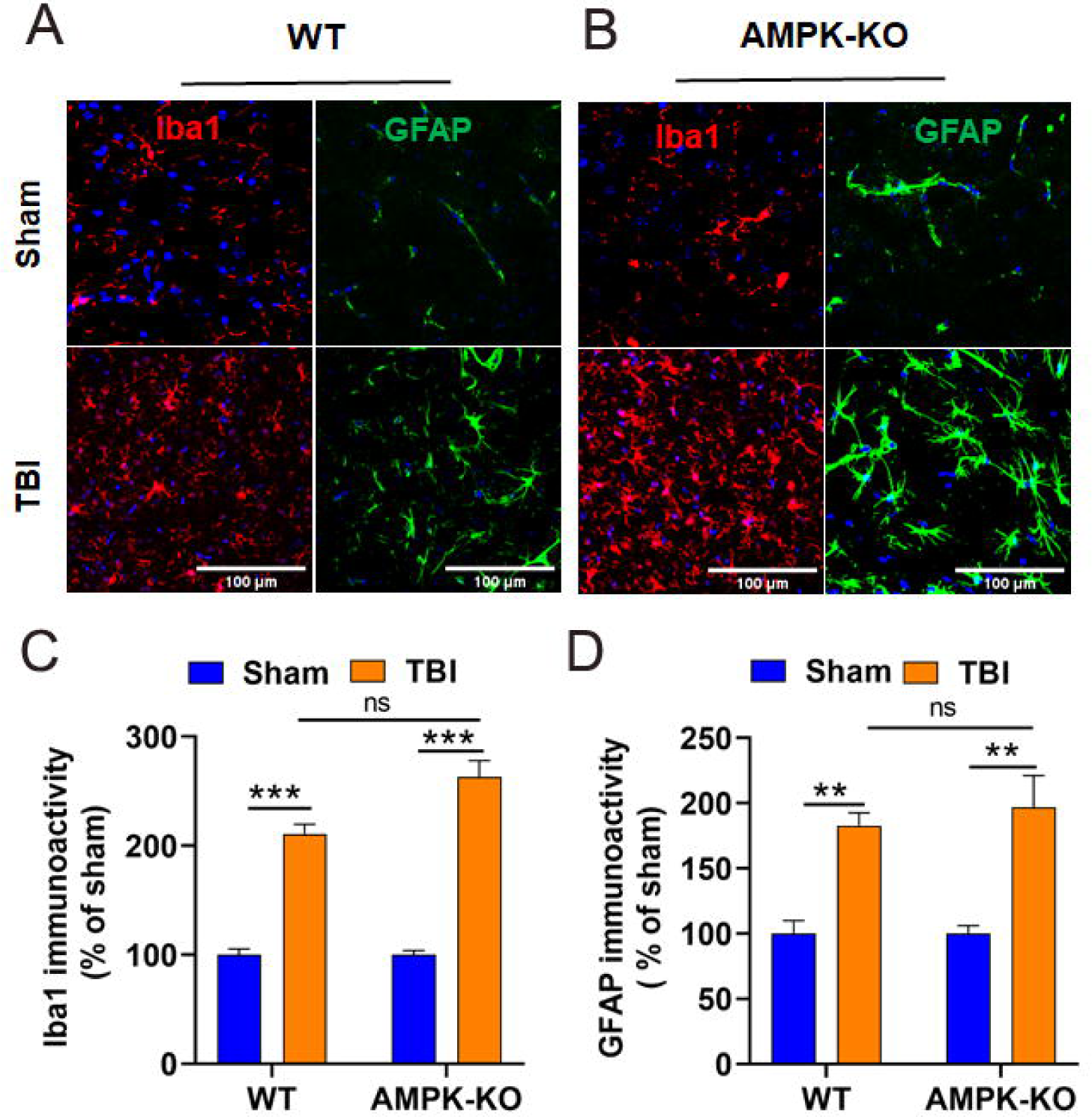
Effect of the absence of AMPK on reactive astrogliosis following TBI. The numbers of astrocytes and microglia in the coronal sections of WT and AMPK-KO mice were determined by immunostaining with anti-GFAP and anti-Iba1 antibodies, respectively. **(A, B)** representative confocal images of brain sections from WT and AMPK-KO mice showing greater expression of GFAP (green) and Iba1 (red) in AMPK-KO mice than in WT mice at 24 h after TBI. Scale bar = 100 µm. **(C, D)** Quantitative analysis of GFAP and Iba1 in the brains of WT and AMPK-KO mice. The data are expressed as 4-5 sections (n=6 mice per group). **p<0.01 *** P<0.001, ns= not significant,

## Discussion

Our study provides novel insight into the role of AMPK in the context of TBI and inflammation. We observed a significant decrease in AMPK activity, as shown by reduced phosphorylation at Thr172 following CCI-induced TBI in mice. This decrease in AMPK phosphorylation coincides with worsened functional impairments and increases neuroinflammation characterized by upregulation of NLRP3 inflammasome. This study showed the impact of AMPK on brain injury. We demonstrated that reduced AMPK levels exacerbate inflammation following TBI. This exacerbation was associated with increased brain tissue damage, heightened neuronal death, and exacerbated functional deficits. Our findings highlight a previously unexplored role for AMPK in modulating inflammation and brain damage following TBI.

The brain undergoes a distinct metabolic response following TBI and immediately after brain injury, there is an increase in metabolic activity characterized by increased glucose utilization lasting up to 30 minutes ^5^. However, this initial hypermetabolic phase is followed by a prolonged period of hypometabolism, which is detectable as early as 6 hours postinjury and persists for days to weeks thereafter ^4,5,50^. In a mouse model of brain injury, activation of the NLRP3 inflammasome has been shown in previous studies to increase inflammation and worsen brain injury ^24,51,52^. In our investigation, we initially examined the phosphorylation status of AMP-activated protein kinase (AMPK) in the ipsilateral cortex 24 hours post-TBI in wild-type (WT) mice. Our results revealed that CCI injury led to reduced AMPK activity, as indicated by decreased phosphorylation at Thr172, as determined by western blot analysis. To further confirm the involvement of AMPK in TBI, we utilized AMPK-KO mice and confirmed that AMPK deficiency exacerbates brain injury. This finding underscores the critical role of AMPK in modulating the pathological processes associated with TBI. Several studies ^53–55^ have emphasized the inflammatory response as a critical factor and primary contributor to complications such as neuronal loss and long-term behavioral deficits following TBI. Previous research has indicated that AMPK plays a key role in reducing the expression of the inflammasome, thereby attenuating brain damage ^56–58^. These findings suggest that the modulation of inflammation by AMPK could be a key mechanism for mitigating TBI-induced neuroinflammation and associated neurological consequences. In our study, as detailed in the results section, we observed that following TBI, the loss of AMPK led to the activation of key inflammatory components, including NLRP3, caspase-1, IL-1β, and ASC, within 24 h post-injury. These findings highlight the therapeutic potential of targeting AMPK to mitigate post-TBI inflammation and associated neurological damage. Our research has highlighted the detrimental impact of AMPK loss on the prognosis of TBI patients.

However, our study has several limitations that merit consideration. First, we focused exclusively on assessing the inflammatory role of AMPK loss using a preclinical mouse model of TBI. We did not explore the potential therapeutic benefits of AMPK activation following TBI in mice. Second, our study primarily demonstrated a reduction in AMPK and subsequent activation of the NLRP3 inflammasome 24 h after TBI. Additional investigations are needed to elucidate the broader temporal dynamics and functional consequences of AMPK modulation in the context of TBI. Therefore, it will be essential to investigate the long-term effect of TBI on AMPK and explore the potential therapeutic benefits of selectively activating AMPK using AMPK activators to further elucidate the mechanism involved in AMPK loss and NLRP3 inflammasome activation post-TBI. These investigations could provide valuable insight into developing targeted therapies aimed at modulating AMPK activity to improve outcomes following TBI.

### Conclusion

Our findings support the hypothesis that the loss of AMPK following TBI is correlated with worsened behavioral outcomes and heightened inflammasome activation. These results suggest that targeting AMPK activation could be a valuable approach for developing treatments for TBI-associated inflammation and improving post-injury outcomes. Consequently, in future studies, the use of a selective AMPK activator will represent a promising therapeutic strategy to mitigate inflammation in the aftermath of TBI.

## Acknowledgments

This work was partially supported by the Department of Neurology Henry Ford Health and NIH award number R21NS123531 to ASA and the National Institutes of Health (NS112727, AI144004) and Henry Ford Hospital Internal support (A10270, A30967) to SG. The funders had no role in study design, data collection, and interpretation, or the decision to submit the work for publication.

